# A model, mixed-species urinary catheter biofilm derived from spinal cord injury patients

**DOI:** 10.1101/2024.11.04.621968

**Authors:** Parisa Noorian, Kailey Hamann, M. Mozammel Hoque, Gustavo Espinoza-Vergara, Joyce To, Dominic Leo, Priyadarshini Chari, Gerard Weber, Obaydullah Marial, Julie Pryor, Iain G. Duggin, Bonsan Bonne Lee, Scott A. Rice, Diane McDougald

## Abstract

Complex multispecies biofilms consistently colonise the interior of indwelling urinary catheters, causing persistent asymptomatic bacteriuria and frequent symptomatic episodes in long-term catheterized individuals. Simple single-species models often fail to capture the complexities of mixed-species interactions, leading to limited conclusions about microbial behaviour and treatment efficacy. Additionally, using lab-based organisms can obscure the genomic diversity found in real-world infections. The primary objective of this study was to establish a stable and reproducible *in vitro* biofilm model derived from the multi-species clinical flora associated with catheter-related infections, reflecting the dynamics of *in vivo* infections. Biofilm samples from clinical catheters of spinal cord injury (SCI) participants were used to establish polymicrobial macro-fluidic models within catheters. Metagenomic techniques using short-read Illumina and long-read Oxford Nanopore sequencing was used to assess the community composition, produce metagenome-assembled genomes (MAGs), analyse strain-level phylogeny diversity and single nucleotide polymorphisms (SNP) of isolates. Antibiotic resistance tests using our models highlighted the drastic differences between planktonic bacteria, single-species, and multispecies biofilms. *In silico* analysis of antibiotic resistance further revealed a high number of varied resistance genes present in these communities. The models developed and characterised in this study are expected to facilitate more effective strategies to prevent and treat catheter-associated infections.

## Introduction

Urinary catheters are commonly used when the bladder cannot empty naturally. The insertion of a catheter provides an abiotic surface free of the host’s innate defence system and is a suitable niche that bacteria can colonise for biofilm formation. This often leads to catheter associated urinary tract infections (CAUTI), which are hard to diagnose and even harder to treat ^1^. CAUTI have potentially life-threatening consequences in part due to selection for antibiotic resistant strains ^2,3^. Biofilms are surface-associated communities, embedded in extracellular polymeric substances (EPS) ^4^. Biofilms are made up of single- or multi-species consortia and can consist of a mixture of Gram-negative and Gram-positive bacteria as well as other microorganisms such as yeasts. Increasing the duration of catheterization is linked to an increased likelihood of biofilm formation, symptomatic infection and increased diversity of the bladder flora ^5^. Examination of cross sections of catheters from patients revealed that biofilms varied in thickness from patchy monolayers of cells to extensive biofilm hundreds of cells deep where the thickness varies from 3 µm to 0.5 mm. In some cases, they extended the whole length of the urinary catheter and with bacterial populations of up to 5 × 10^9^ colony-forming units (CFU) cm^-2^ of luminal surface area ^6^. Biofilms, especially mixed species communities, are particularly challenging to treat compared to planktonic cells or single species biofilms. Multi-species bacterial communities exhibit social behaviours including changes in virulence, production of secondary metabolites, horizontal gene transfer, colonisation potential and persistence, biofilm development and increased stress resistance, e.g. antimicrobial compound resistance ^7,8^.

To effectively study and combat these complex biofilms, it is important to develop more sophisticated models that accurately reflect the dynamics of *in vivo* infections. Simple single or dual species models often fail to capture the intricacies of mixed-species interactions. A number of different bacterial biofilms models have been developed to better understand how biofilms form and how to eradicate them. Such studies have shown varying outcomes depending on the tested isolates as well as the surfaces or media used for testing. For example, dual species biofilms grown on silicon coupons immersed in artificial urine medium showed that *Escherichia coli* cell numbers decreased when co-cultured with *Pseudomonas aeruginosa*, while *P. aeruginosa* seemed to benefit in those dual-species biofilms ^9^. In contrast, pre-colonisation of catheters by cocultures of *E. coli* with non-pathogenic species, such as *Delftia tsuruhatensis* and *Achromobacter xylosoxidans*, within dual-species biofilms promoted adhesion of *E. coli* ^10^.

Another study comparing single and dual species biofilms of isolates originated from catheter-associated polymicrobial bacteriuria on siliconized coverslips in artificial urine showed *Klebsiella pneumoniae* formed biofilms in co-incubation with *E. coli* or *Enterococcus faecalis*, but co-inoculation with *P. mirabilis* impaired its biofilm formation. *E. coli* did not persist in biofilms during co-incubation with *K. pneumoniae*, but was not affected by *E. faecalis*. *E. faecalis* surface attachment was partially inhibited by *E. coli*, but not by *K. pneumoniae* ^11^. These observations are expected to be highly strain-dependent, which underscores the necessity of clinically relevant multi-species models, as the genetic background of laboratory strains of bacteria in isolation do not reflect the intricacies and dynamics where multi-species are present.

Biofilm-associated pathogens have a much higher survival rate in comparison with their planktonic counterparts against stresses such as nutrition deprivation, extremes in salinity, pH, temperature, UV, chlorine, antimicrobial agents and phagocytosis by immune cells by one or more mechanisms. Multispecies biofilms have an even a higher rate of resistance to various stresses compared to single-species biofilms, and both types of biofilms are significantly more resistant compared to planktonic bacteria ^12-14^. When interventions alter the balance of microbial communities, they can influence competitive interactions, resource availability, and niche occupation. For example, a static, synthetic multi-species biofilm showed that low nutrient conditions and sub-lethal concentrations of an alcohol-based disinfectant enhanced biofilm biomass. Furthermore, a reduction of the dominant taxa in the multi-species biofilm as a result of the sub-lethal disinfectant concentrations lead to an increase in antimicrobial-resistant minor taxa ^15^.

Various static, microwell and microfluidic-based biofilm formation systems are widely used. Each system has benefits and disadvantages but regardless of the system, *in vitro* biofilm models often show high variability between results of different systems, preventing reliable comparison of treated biofilms to their controls ^16,17^. To design an *in vitro* model representative of real-life patients, there are many important factors to be considered, including the substratum material, growth conditions, microbial community, as well as the physiological and immunological responses of the host ^18^. This discrepancy in models arises due to differences in surface properties, the absence of biological interactions, surface uniformity, variations in hydrophobicity and hydrophilicity, lack of shear forces, insufficient antimicrobial defences, and altered biofilm architecture. Due to these challenges, it is impractical to control all aspects of the bladder environment in a single model.

Our approach focuses on achieving consistent outcomes under the flow of artificial urine inside a urinary catheter using clinically relevant bacterial isolates. For this purpose, bacterial communities isolated from spinal cord injury (SCI) participants were used to establish a repeatable multispecies biofilm on the intraluminal surface of catheters. Here, we used synthetic urine medium, which imitates nutrient conditions within the bladder better than nutrient rich media, and can be consistently produced in large volumes required for macro-fluidic systems. High throughput sequencing and metagenomic analysis were used to study the development of the community composition at each step as well as to identify the specific genes, e.g. antimicrobial genes and virulence genes, associated with those community members.

Here, we also used our models to evaluate the efficacy of existing treatments against complex biofilm-associated infections, providing valuable data to improve clinical practices. Our multispecies biofilms are relevant to clinical bacterial samples, and can serve as a robust platform for experiments with novel antimicrobial compounds, biofilm-disrupting agents, and alternative therapeutic strategies in a controlled and reproducible environment. This would further allow us to investigate the basic principles of microbial interactions, biofilm formation, and gene expression within a clinically relevant context.

## Results

### Establishment of an *in vitro* multi-species biofilm from the bacterial community of an SCI patient

In this study, our main goal was to establish a stable, reproducible, *in vitro* multi-species biofilm community on an intraluminal catheter derived from clinical catheter-associated flora and closely define the community’s features. For this purpose, bacterial communities from the indwelling catheters of six SCI patients with neurogenic bladders were selected. The participants were at least five years post-initial injury and at least 2 months post last symptomatic CAUTI. All samples were explanted as part of the participant’s routine catheter change and were considered asymptomatic bacteriuria (Table 1).

**Table 1.** Details of participants and the catheters at the time of explantation.

| Model | Gender | Age | Years since SC injury | Months since last CAUTI | Weeks since last catheter change | Catheter type | Catheter size and brand |
| --- | --- | --- | --- | --- | --- | --- | --- |
| C-1 | Male | 65 | 11 | 3 | 4 | Suprapubic | Mdevices 18 G |
| C-2 | Female | 44 | 43 | >3 | 3 | Suprapubic | Releen 18G |
| C-3 | Male | 74 | 56 | >3 | 4 | Suprapubic | Releen 18G |
| C-4 | Female | 35 | 5 | 2 | 3 | Suprapubic | Uromed double balloon 18G |
| C-5 | Female | 72 | 47 | >6 | 4 | Suprapubic | Releen 18G |
| C-6 | Male | 66 | 5 | >6 | 4 | Suprapubic | Uromed double balloon 20G |

The biofilm from each catheter was extracted immediately after they were received and sequenced using metagenomic shotgun sequencing to determine the bacterial community population at the time of explantation. Analysis using Kraken2 showed that catheters (C-1 to C-6) were colonised by a range of bacterial species. In total, 87 bacterial species were found across the 6 samples (Figure 1). A portion of the biofilm was stored in 25% glycerol at -80°C and were later grown planktonically in complete composite Synthetic Human Urine (ccSHU)^19^ for 48 h at 37°C under static conditions for subsequent use as input (“In”) for biofilm formation. The biofilm was cultivated in an intralaminar urinary catheter using a macro-fluidic-based flow system and extracted after 4 d to be used as output (“Out”). This process was repeated 3 consecutive times (G1 to G3), each time using output biofilm to seed the next input. To ensure the reliability and reproducibility of our results, we conducted the whole experiment with three independent replicates. At the same time, bacterial diversity was also assessed on CHROMagar™ Orientation plates based on colony colour and morphology.

**Figure 1.**
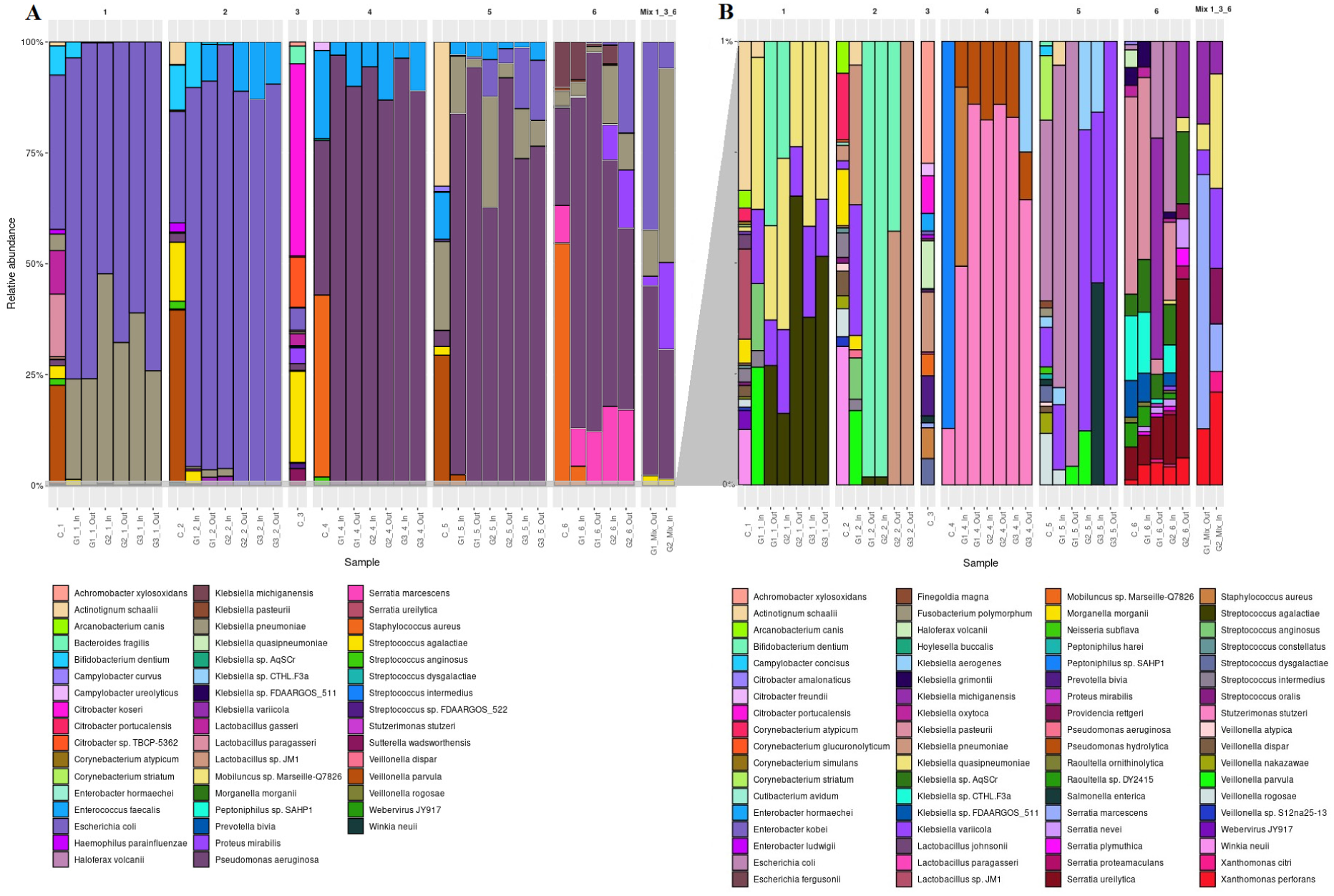
Metagenomic reconstructions of community composition and representation by species. Relative abundance plots of catheter biofilms species are plotted. A) High abundance bacteria ≥ 1% to 100%. B) Less abundant bacteria, <1% of total reads. The bacterial abundance of catheters explanted from participants are labelled C-1 to C-6. The bacterial abundance of catheters going through 3 consecutive generations of the flow model are labelled G1 to G3, biofilm model number and “In” for input planktonic inoculation bacteria or “Out” for output biofilm bacteria.

C-1 was mainly colonised by E. coli (31.95%), Veillonella parvula (20.42%), Lactobacillus paragasseri (13%), Lactobacillus gasseri (9%), Bifidobacterium dentium (6%), K. pneumoniae (3.41%), Streptococcus agalactiae (2.69%), Streptococcus anginosus (1.36%), P. aeruginosa (1.21%) and Haemophilus parainfluenzae (1%). The normally aerotolerant anaerobes or microaerophilic Lactobacillus spp. and B. dentium, which originally compromised 28% of the total reads, did not thrive in the input cultures under the conditions used (Figure 1). Interestingly, P. aeruginosa and H. parainfluenzae were also completely lost from first in vitro Input growth, while V. parvula and B. dentium could not be detected after the first (G1_1_Out) and second (G1_2_Out) passages of biofilm formation, respectively. The final community abundance of the C-1 biofilm model shifted to E. coli (71.77%) and K. pneumoniae (24.81%) at higher abundance (>1% of reads) and S. agalactiae (0.16%), K. quasipneumoniae (0.11%) and K. variicola (0.04%) at lower abundance (<1% of reads). E. coli and Klebsiella spp. were identified on CHROMagar™ Orientation plates and Even though S. agalactiae was at low abundance, 0.16% of the community, it was readily recovered.

C-2 was mainly colonised by 9 bacterial species, *V. parvula* (38.37%)*, E. coli* (24.89%)*, S. agalactiae* (13.15%)*, B. dentium* (10 %)*, Actinotignum schaalii* (5%)*, P. aeruginosa* (2%)*, H. parainfluenzae* (2%) and *S. anginosus* (1%). After 3 generations of biofilm formation, the community abundance shifted so that only *E. coli* and *E. faecalis* remained at 90.32% and 9.50% of the total reads, respectively. *E. faecalis* were recovered on CHROMagar™ Orientation plates.

C-3 was colonised by at least 10 bacterial species (>1% of reads), *Citrobacter koseri* (43%)*, S. agalactiae (20.43%), Citrobacter* sp. TBCP-5362 (11.28%)*, E. coli* (4.95%)*, Bacteroides fragilis* (3.97%)*, Sutterella wadsworthensis* (3.79%)*, P. mirabilis* (3.55%)*, L. gasseri* (2.69%)*, P. aeruginosa* (1.51%) and *Streptococcus sp. FDAARGOS_522* (1.22%). No biofilm was formed from inoculation of this community and no amount of DNA was recovered from the *in vitro* cultures.

C-4 was colonised mainly by *Staphylococcus aureus* (40.81%)*, P. aeruginosa* (34.62%)*, E. faecalis* (19.73%)*, Campylobacter ureolyticus* (1.93%) and *S. anginosus* (1.91%). The community abundance of C-4 changed to *P. aeruginosa* and *E. faecalis* after the first *in vitro* growth. *S. aureus* which was more than 40% of the reads in C-4 decreased to under 1% in first Input cultures and could not be detected in subsequent passages. Whereas *P. aeruginosa* and *E. faecalis* compromised 88.34% and 11.01% of the total reads, respectively. *E. faecalis* and *P. aeruginosa* were recovered on CHROMagar™ Orientation plates.

C-5 was colonised mainly by *Actinotignum schaalii* (32.4%), *V. parvula* (29.16%)*, K. pneumoniae* (19.94%)*, E. faecalis* (10.59%)*, P. aeruginosa* (3.55%)*, S. agalactiae* (1.95%) and *Campylobacter curvus* (1.18%). After *in vitro* growth, the community shifted to *P. aeruginosa* (76.48 %), *E. coli* (13.53%), *K. pneumoniae* (5.82%) and *E. faecalis* (4.13%). On the original catheter *E. coli* was only 0.46% of the reads and reached 13.53% of the reads by G3_5_Out. *P. aeruginosa* also increased from 3.55% in the patient’s catheter sample to 76.48% of the reads in G3_5_Out. *K. pneumoniae, E. faecalis, P. aeruginosa* and *E. coli* were recovered on CHROMagar™ Orientation plates.

C-6 was colonised by mainly *S. aureus* (54.28%)*, P. aeruginosa* (21.87%)*, Klebsiella michiganensis* (10.22%)*, Serratia marcescens* (8.43%) and *K. pneumoniae* (3.33%). The biofilm extracted from the second generation of biofilm formation (G2-6-out) showed that the community shifted to *P. aeruginosa* (40.78%), *E. coli* (20.48%)*, S. marcescens* (16.89%)*, P. mirabilis* (13.1%) and *K. pneumoniae* (8.28%)*. P. mirabilis* was initially detected as 0.00053% on the patient’s catheter but increased to 13.1% by G2-6-out. Similarly, *E. coli* was found at 0.02% and rose to 20.48% by G2-6-out. The generation 3 planktonic input (G3-6-In) did not grow in ccSHU and no detectable DNA was recovered.

Models C-1, C-3 and C-6 were mixed at a ratio of 1:1:1 (G1_Mix_Out and G2_Mix_In) containing *K. pneumoniae* (43.62%), *P. aeruginosa* (29.14%), *P. mirabilis* (19.53%), *E. coli* (5.95%) and *S. agalactiae* (1.45%). No biofilm was formed from inoculation of the G2_Mix_In and no DNA was recovered.

In summary, biofilms of models C-1, C-2 and C-4 developed *in vitro* were primarily compromised of two major species, while biofilm of model C-5 had four major species. Whereas, biofilms of models C-3, C-6 and C-1, C-3, C-6 mixture could not be established in the laboratory. The reasons for the failure to establish these biofilms remain unclear.

### Strain diversity across models

Separating different strains of the same species in metagenomic analyses presents significant challenges due to the high degree of genetic similarity and the complexity of microbial communities within strains. To determine differences in the dominant strains between models and potential changes within each passage, we utilised the StrainPhlAn pipeline with its default settings, which selects the allele with more than 80% dominance ^20^. This analysis focused on the four prominent species found in our models: *E. coli*, *K. pneumoniae*, *E. faecalis*, and *P. aeruginosa*. In all of the models, the strains remained consistent through *in vitro* passages and group together on the phylogenetic tree.

*E. faecalis* strains found in models C-2, C-4, and C-5 are phylogenetically distinct from one another, with none showing close genetic relationships to the reference strains that were tested. In contrast, the *E. coli* strains found in C-1 were distinct from those in models C-2 and C-5. C2 and C5 participants were recruited from the same enrolment site. Notably, the *E. coli* strain in model C-1 is close to UPEC reference strains UTI89 and urine serovar O1:K1:H7 type strain DSM 30083 (ATCC 11775). Screening of the *E. coli* O-groups and H-types (EcOH) database using the Abricate pipeline revealed that the *E. coli* strain in model C-1 belongs to serotype O2:H1 while those in model C-5 belong to serotype O18:H5.

Additionally, the *K. pneumoniae* strain in model C-5 differs from models C-1 and C-6 and is more closely aligned with the tested reference genomes. In terms of phylogenetic proximity, the strains from model C-1 and C-6 and as a result the mixed model demonstrate similar relationships on the phylogenetic tree. Furthermore, the *P. aeruginosa* strains found in models C-4, C-5 and C-6 are distinct from one another. Model C-6 dominant strain is closest to the reference genome *P. aeruginosa* PAC-1 which is recognized for its high levels of multidrug resistance.

### Isolation of members of each community

Based on final diversity in the *in vitro* system, models C-1 (2 major species) and C-5 (4 major species) were selected for further analysis. G3_1_Out and G3_5_Out were used as inoculants for the Input (G4) cultures used for all subsequent experiments. These cultures were sequenced using Oxford Nanopore Technology (ONT) ultradeep long read metagenome sequencing on a nanopore PromethION as well as short read Illumina shotgun sequencing. Hybrid short and long read assembly was used for generating metagenome-assembled genomes (MAGs). Both sets of MAGs were of high quality: >90% complete, <5% contaminated (Supplementary Data 1). In model C-1, *E. coli* and *K. pneumoniae* were completed at 100% and *S. agalactiae* was completed at 92.49%. For model C-5, *K. pneumoniae, E. faecalis* and *E. coli* were completed at 100% and *P. aeruginosa* was completed at 96.35%, as calculated by CkeckM2. GTDB-Tk results confirmed the species assignment.

Input (G4) cultures of models C-1 and C-5 were also spread on CHROMagar™ Orientation plates at each passage. Three colonies of each colour and morphology were selected and short read shotgun sequencing was used to assemble genome of each isolate. The taxonomic affiliation of the genomes was determined using GTDB-Tk v2.4. One isolate from each phenotype is presented in Table 2, providing a representative overview of the genetic diversity and taxonomic classification of single isolates within the samples.

**Table 2.** Summary of single isolates from models C-1 and C-5 community selected based on phenotypic differences.

|  | Isolate | Colony colour* | Reference genome | skani_ani | skani_af | Reference taxonomy |
| --- | --- | --- | --- | --- | --- | --- |
| Model C-1 | 1-1 | Light Blue | GCF_000186445.1 | 98.54 | 0.8593 | *Streptococcus agalactiae* |
|  | 1-2 | Pink | GCF_003697165.2 | 98.50 | 0.8999 | *Escherichia coli* |
|  | 1-5 | Metallic blue with pink halo | GCF_000742135.1 | 98.90 | 0.8855 | *Klebsiella pneumoniae* |
| Model C-5 | 5-1 | Pink | GCF_003697165.2 | 98.53 | 0.9004 | *Escherichia coli* |
|  | 5-3 | Turquoise blue | GCF_029024925.1 | 98.87 | 0.8788 | *Enterococcus faecalis* |
|  | 5-5 | Metallic blue no halo | GCF_000742135.1 | 99.03 | 0.8847 | *Klebsiella pneumoniae* |
|  | 5-6 | Light Green | GCF_001457615.1 | 99.39 | 0.9607 | *Pseudomonas aeruginosa* |
|  | 5-10 | Cream | GCF_001457615.1 | 99.45 | 0.9572 | *Pseudomonas aeruginosa* |

\* Colony colour based on pigment production on CHROMagar™ Orientation plates.

There were two morphotypes of *P. aeruginosa*, based on pigment: one producing a yellow-green pigment (strain 5-6) and the other non-pigmented (strain 5-10) (Table 3). Genomic comparisons of both *P. aeruginosa* isolates revealed no transposons or large genomic insertions. To identify genetic changes underlying the altered phenotypes, we used single nucleotide polymorphism (SNP) analysis with Breseq v0.38.3 to compare the genomic sequences of isolates from each morphotype. Specifically, three yellow-green pigmented isolates were compared to three non-pigmented isolates, considering only mutations present in 100% of sequencing reads. The analysis identified 38 non-synonymous mutations, 3 synonymous mutations, and 4 insertions and deletions (INDELS), with a total of 25 hypothetical proteins found to have SNPs. None of the genes carrying SNPs were directly associated with biosynthesis of pigments.

**Table 3.**
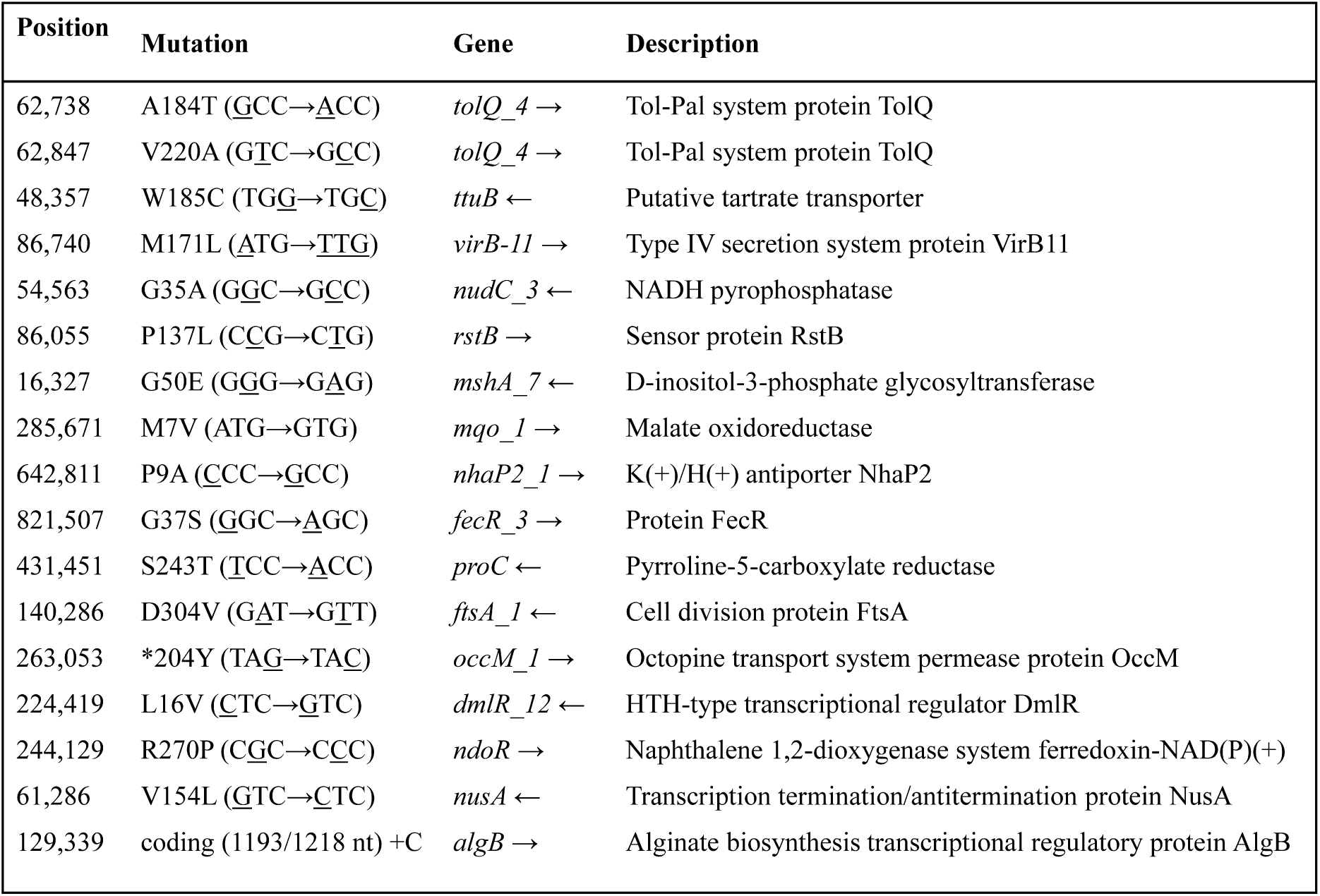
Selected SNP differences between *P. aeruginosa* isolated with two distinct phenotypes (yellow-green pigmented (strain 5-6) or non-pigmented (strain 5-10)) isolated from model C-5. Three isolates from each phenotype were selected for comparison and only mutations present in 100% of the sequencing reads are reported. Underlined bases denote single substitution.

### Metagenomic analysis of antibiotic resistance mechanisms

We utilised the metgenomic sequences to investigate the mechanisms of antimicrobial resistance and to determine if the microorganisms carried any additional resistance mechanisms or genetic markers. We employed Abricate to scan each MAG against Comprehensive Antibiotic Resistance Database (CARD) and NCBI AMRFinderPlus in order to detect AMR genes. The single isolates were also analysed in similar fashion and crossed checked with metagenomics analysis.

This analysis yielded a total of 7 matches from NCBI and 57 matches from CARD for model C-1 while model C-5 produced an alarming 23 matches from NCBI and 115 matches from CARD (Figure 3). The identified ARGs from screening of CARD were classified into two categories. The first category, “Efflux Systems,” includes two-component regulatory systems modulating antibiotic efflux and efflux pump complex or subunit associated with antibiotic resistance in the CARD database. The second category, labelled “Other,” encompasses all remaining ARGs, such as enzymes, porins, and peptides (Supplementary Data 2). These two clinical models have a large range of antibiotic resitance that can be categorized into twenty families: aminoglycosides, cephalosporin, fluoroquinolones, glycylcyclines, peptides, phenicols, rifamycins, fosfomycins, aminocoumarins, nitroimidazoles, diaminopyrimidines, macrolides, sulfonamides, tetracyclines, lincosamides, triclosan, benzalkonium chloride, acridine dye and nucleosides.

**Figure 2.**
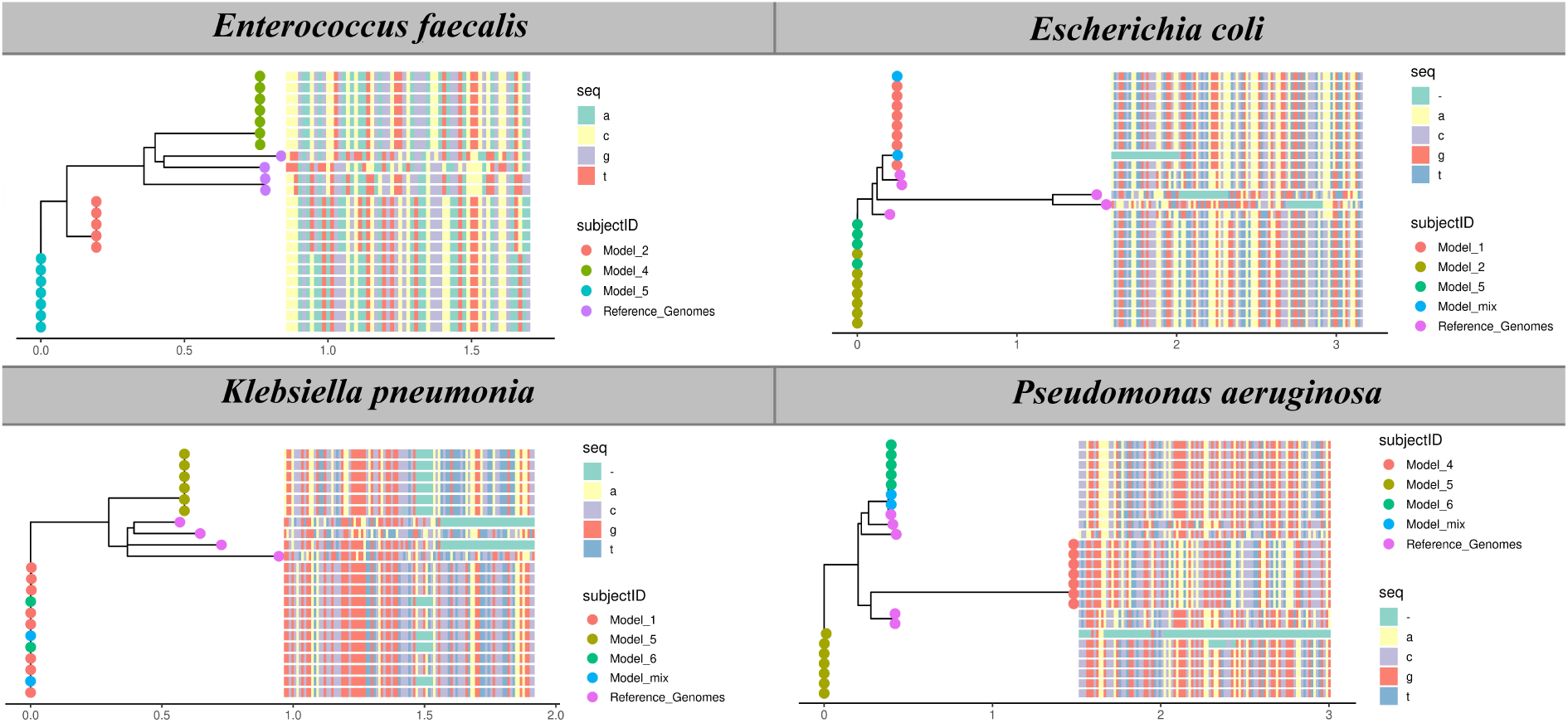
Maximum likelihood phylogenetic tree of the four major species profiled in the models. Species profiled using StrainPhlAn are *E. coli*, *K. pneumoniae*, *E. faecalis*, and *P. aeruginosa*. The branch tips of the phylogenetic tree are color-coded according to the different models used. Accompanying the phylogenetic tree is a slice of the multiple sequence alignment that highlights unique gene marker sequences.

**Figure 3.**
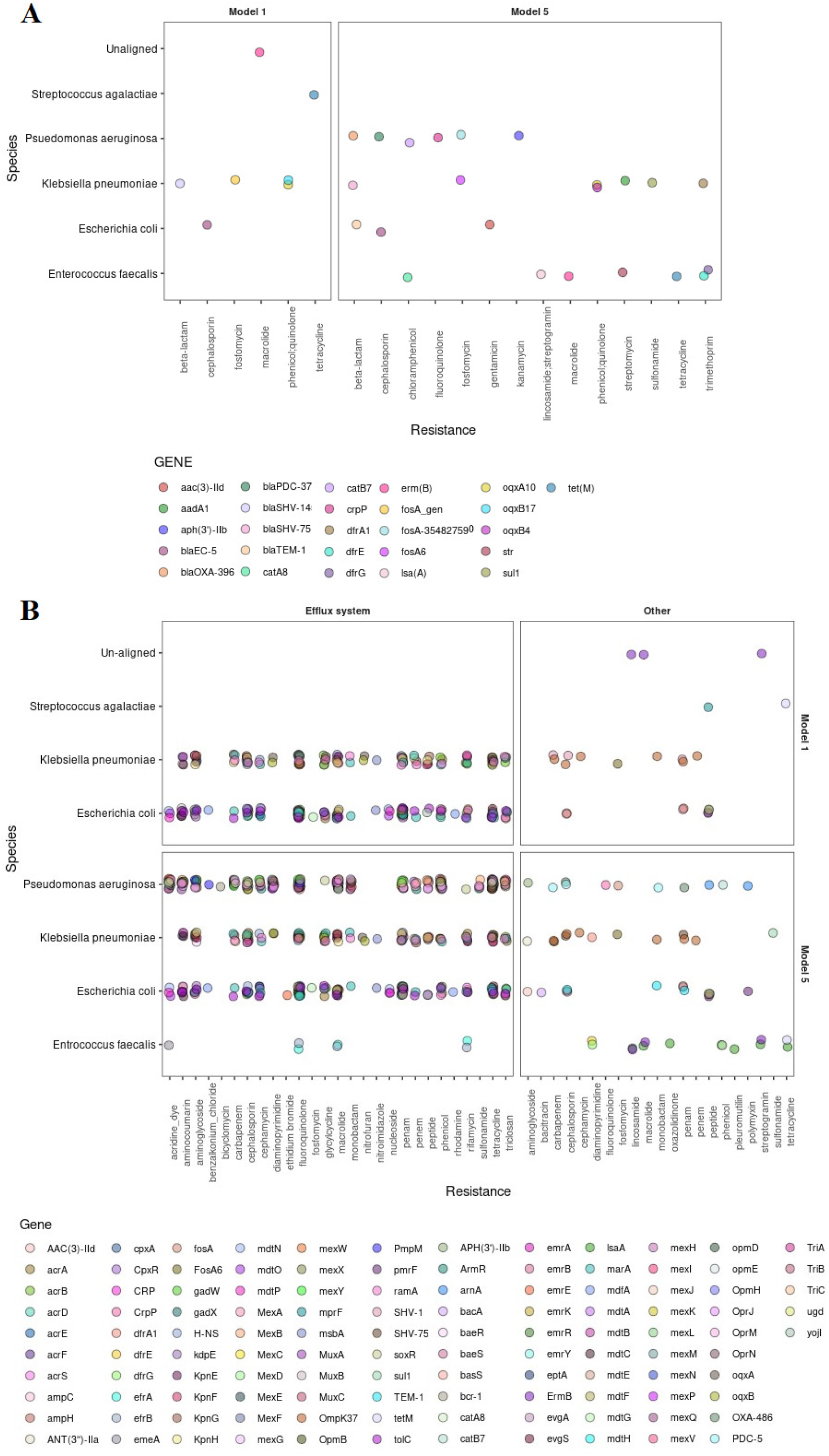
Summary overview of A) NCBI AMRFinderPlus and B) CARD show the ARGs detected in MAGs of model C-1 and model C-5.

From model C-1, *S. agalactiae* had the lowest number matches (three) to antibiotic resistance genes for all of the databases used here, *tetM* (tetracycline), *ErmB* (erythromycin) and *mprF* (cationic peptides). In contrast, *E. coli* had 43 matches against the CARD database. Similarly, *K. pneumoniae* had 22 matches against CARD.

In model C-5, *E. faecalis* strain 5-3 showed 10 matches in CARD. *E. coli* from model C-5 had 45 matches, while *K. pneumoniae* had 25 matches. *P. aeruginosa* also showed a significant number of potential resistance genes, with 47 matches in CARD, and no differences were observed when the isolates were individually assessed between the individual isolates 5-6 and 5-10.

The screening of NCBI revealed no genes specifically associated with gentamicin resistance in model C-1. In contrast, in model C-5, the *E. coli* strain carried the *aac(3)-IId* gene, potentially conferring direct resistance to gentamicin (Figure 3-A). Further investigation through CARD revealed a notable presence of genes potentially involved in aminoglycoside resistance. Specifically, 13 genes in model C-1 and 22 such genes model C-5 (Figure 3-B). All of the predicted genes in both models were related to efflux pump systems. Four efflux pump-related genes, *acrD, baeR, cpxA* and *KpnE* were detected in *E. coli* and *K. pneumoniae* from both models. Additionally, the *tolC* was detected in *E. coli* in both models. In *K. pneumoniae* from both models *kpnF, kpnG,* and *kpnH* were present. Furthermore, *emrE, pmpM* genes and complex *mexXY-oprM* genes were detected in *P. aeruginosa* in model C-5 (Supplementary Data 2). Interestingly, *S. agalactiae* from model C-1 and *E. faecalis* from model C-5 did not show any genes previously associated with gentamicin resistance.

### Antibiotic resistance profile of bacterial isolates

Antibiotic resistance is a crucial phenotype for understanding both bacterial communities and their individual members in a catheterized patient population. Here, we profiled antibiotic resistance of single isolates from model C-1 and C-5 by utilising a standard Kirby-Bauer disk diffusion test to evaluate the effectiveness of 22 antibiotics. The antibiotics tested belong to several families including aminoglycosides (amikacin, gentamicin, kanamycin, neomycin, netilmicin, spectinomycin, streptomycin, tobramycin), Beta-lactams including penicillins (ampicillin sulbactam, ampicillin), cephalosporins (cefotaxime, ceftazidime, ceftriaxone), carbapenems (imipenem and meropenem), chloramphenicol, Fluoroquinolones (ciprofloxacin and nalidixic), sulfonamides (sulfamethoxazole and trimethoprim), tetracycline and rifampicin (Table 4 A). The diameter of zone of inhibition is measured to determine the susceptibility of the bacteria to the antibiotic. Larger zones of inhibition indicate greater sensitivity, whereas smaller zones suggest resistance. The interpretive standards for each species is used for the determination of antibiotic sensitivity ^21,22^.

**Table 4.**
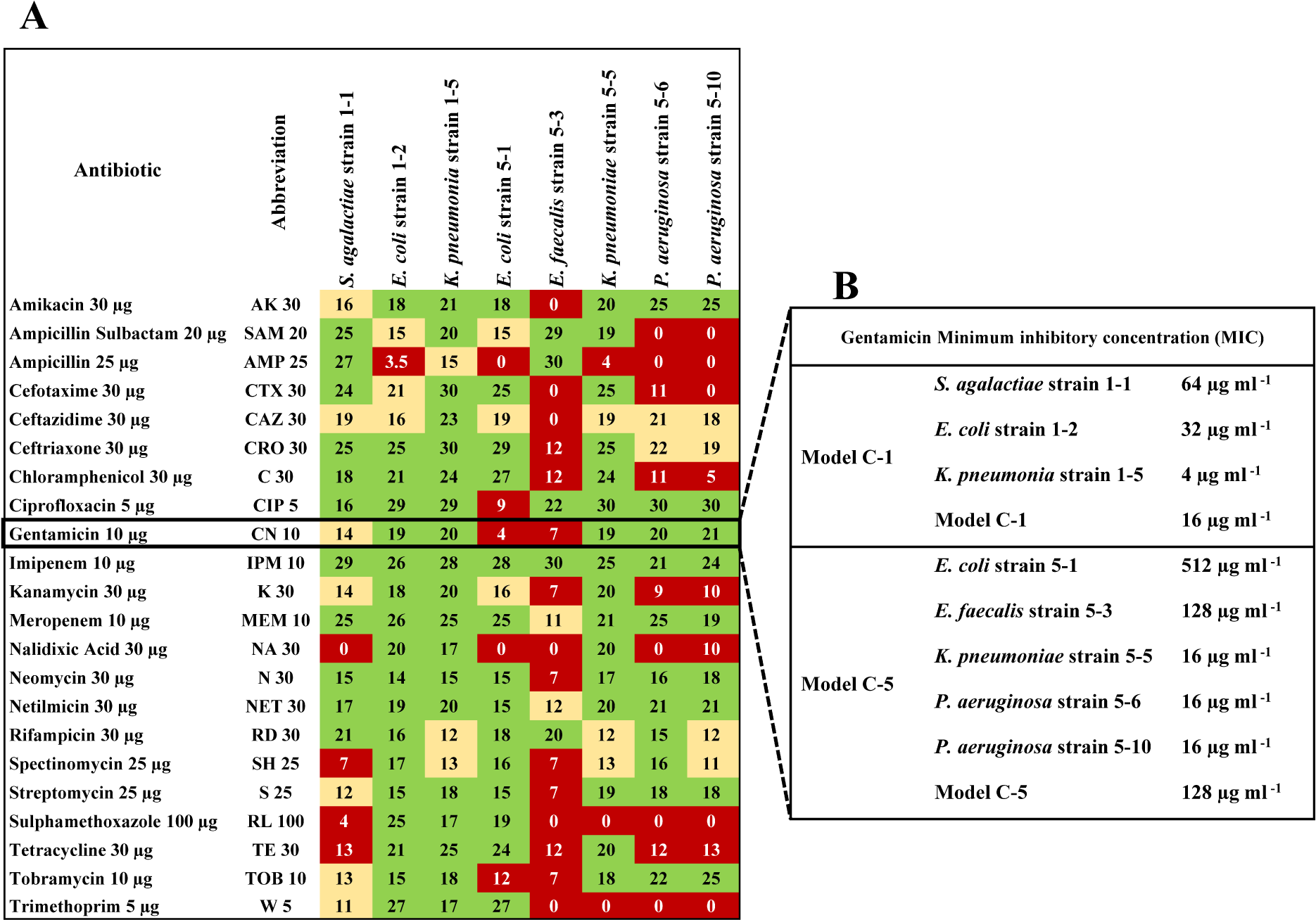
(A) Zone of inhibition of antibiotic disk diffusion for each isolate is reported in diameter (mm). The interpretive standards for the determination of antibiotic sensitivity were used ^21,22^ and susceptibility is reported as resistant (red), intermediate (yellow) and sensitive (green). (B) The minimum inhibitory concentration (MIC) of gentamicin in a microdilution assay based on resazurin reduction for individual species and the multi-species bacterial community. All experiments were repeated 3 times separately with three replicates in each independent experiment.

In general, all strains showed resistance to several antibiotics at varying degrees. The only antibiotic to which all isolates were sensitive was the last resort carbapenem antibiotic, imipenem. All isolates except for *E. faecalis* strain 5-3, were sensitive to netilmicin and meropenem. *E. faecalis* strain 5-3 showed the most resistance from all tested isolates and was resistant to 15 of the tested antibiotics and showed intermediate resistance to another 2 (Table 4 A).

The recommended first-line antibiotic drugs of choice regardless of area of involvement in the urinary tract, include ciprofloxacin or sulfamethoxazole/trimethoprim. All isolates from model C-5, except for *E. coli* strain 5-1, were completely resistant to sulfamethoxazole/trimethoprim. While in model C-1 only *S. agalactiae strain* 1-1 showed resistance to sulfamethoxaloze/trimethoprim. In contrast, from all isolates tested, only *E. coli* strain 5-1 was resistant to ciprofloxacin.

*S. agalactiae strain* 1-1 from model C-1 showed intermediate gentamicin resistance, whereas *E. coli* strain 5-1 and *E. faecalis* strain 5-3 from model C-5 were resistant to gentamicin. To further test the resistance against gentamicin, single isolates or the mixed models were grown planktonically in ccSHU and treated with different concentrations of gentamicin in a microdilution assay (Table 4 B). The MICs from the microdilution assay were consistent with the disk assay. *S. agalactiae strain* 1-1 had an MIC of 64 µg ml^-1^ but the *E. coli* 1-2 (32 µg ml^-1^), *K. pneumonia* 1-5 (4 µg ml^-1^) and the mixed model C-1 (16 µg ml^-1^) have very low MICs. In model C-5, *E. coli* strain 5-1 (MIC of 512 µg ml^-1^) and *E. faecalis* strain 5-3 (MIC of 128 µg ml^-1^) were considered to be gentamicin resistant. Whereas, *K. pneumonia* 5-5 and *P. aeruginosa* 5-6 and 5-10 had lower MICs, 16 µg ml^-1^. The mixed model C-5 showed higher resistance, with an MIC of 128 µg ml^-1^.

### Antibiotic resistance in biofilms

Gentamicin is generally used as both intravesical instillations at 4 mg ml^-1^ for 15 min or intramuscular and intravenous injections against CAUTIS ^23^. Following 80 – 120 mg gentamicin IV or IM, the bulk of the gentamicin is excreted in the urine within the first 24 h after administration and varies from 3 to 600 µg ml ^-1^ ^24,25^. Therefore, the effect of gentamicin on biofilms was tested at either 4 mg ml^-1^ for 15 min or at 500 µg ml ^-1^for 24 h. To compare the effectiveness of gentamicin against isolates in biofilms, we established biofilms using two different systems, biofilms grown on catheter pieces fixed to the lid of multi-well plates or under flow intraluminal catheters pieces. To quantify the culturable cells in each biofilm, CHROMagar™ Orientation plates were utilised.

First, we used catheter pieces fixed to the lid of 48 well plate as pegs to establish 2 d biofilms in ccSHU before treatment with 500 µg ml ^-1^ gentamicin for 24 h (Figure 4). Biofilms were tested as either single species or mixed species in triplicates. In model C-1, when evaluated as individual isolates, *S. agalacticae* 1-1 exhibited the highest resistance to 24 h of gentamicin treatment, showing only a one-log reduction from 2.2 × 10^5^ to 3.2 × 10^4^ CFU cm^-2^. In contrast, *E. coli* 1-2 exhibited a three-log decrease, from 2.96 × 10^7^ to 5.2 × 10^4^ CFU cm^-2^ and *K. pneumoniae* 1-5 was completely sensitive and had a biofilm biomass of 1.3 × 10^8^ CFU cm^-2^ without any treatment, while no colonies could be recovered following antibiotic treatment. When model C-1 was grown as mixed species culture, both *E. coli* and *K. pneumoniae* showed resistance against gentamicin treatment with their biomass decreasing less than one log. *S. agalacticae*, which was only around 0.1% of the population based on metagenomic analayis, was rarely observed in the CFU counts of the mixed model biofilm (Figure 4 A).

**Figure 4.**
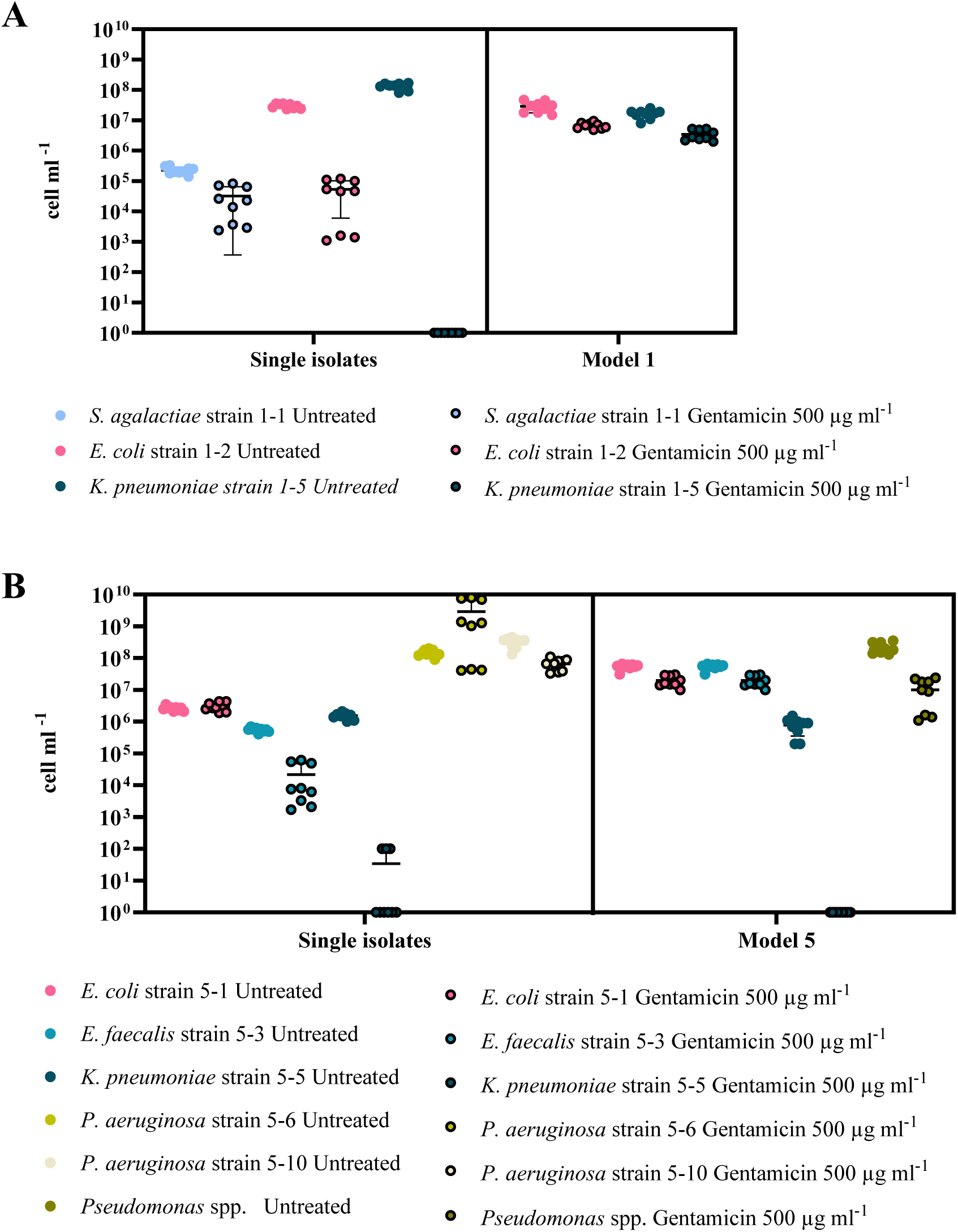
CFU counts on BHI agar plates for single isolates and CHROMagar™ Orientation plates for mixed multi-species models were used to determine the amount of viable culturable bacteria exposed to 500 µg ml ^-1^ gentamicin for 24 h. A) model C-1 and B) model C-5. All experiments were repeated three times separately with three replicates in each independent experiment.

For model C-5, as single species biofilms, *E. coli* 5-1 demonstrated resistance to gentamicin, as anticipated based on the MIC analysis, with 2 × 10^6^ CFU cm^-2^ in untreated biofilms before and after treatment. *E. faecalis* 5-3 was also resistant to gentamicin, albeit to a lesser extent, with a one-log decrease in biomass from 5.5 × 10^5^ to 2.1 × 10^4^ CFU cm^-2^. In contrast, *K. pneumoniae* 5-5 was sensitive to gentamicin, resulting in a five-log decrease from 1.5 × 10^6^ CFU cm^-2^ to 3.4 × 10^1^ CFU cm^-2^ after antibiotic treatment. Both *P. aeruginosa* isolates exhibited gentamicin resistance but with notable diffrences in their response. *P. aeruginosa* 5-6 formed 1.4 × 10^8^ CFU cm^-2^, which increased to 2.9 × 10^9^ while *P. aeruginosa* 5-10 experienced a slight decrease in biomass, reducing from 3.2 × 10^8^ to 6.6 × 10^7^ CFU cm^-2^. When grown as a multi-species biofilm, *E. coli* and *E. faecalis* exhibited no significant changes in their biomass after treatment, with both species maintaining levels around 5 × 10^7^ CFU cm^-2^. Surprisingly, *K. pneumoniae* demonstrated the same sensitivity as when grown as a single isolate, with its biomass completely diminishing from 7.7 × 10^5^ CFU cm^-2^ to no recovarable colonies. *P. aeruginosa* showed a slight decrease, reducing from 2.2 × 10^8^ CFU cm^-2^ to 1.1 × 10^7^ CFU cm^-2^ (Figure 4 B).

Gentamicin resistance of the multi-species biofilm communities grown intraluminal human catheter under flow conditions in ccSHU were tested. To ensure there was no mechanical disruption of the biofilm as a consequence of injection of the treatment, we showed that gentle injection of saline did not disrupt the biofilms and there were no statistical differences in Models C-1 and C-5 biofilm biomass with or without saline injection.

To test the effect of high doses of gentamicin as routinely used as intravesical intervention or catheter irrigation, 4 d biofilms were exposed to 4 mg ml^-1^ for 15 min (Figure 5 A and C). A 15 min treatment of multispecies model C-1 with 4 mg ml^-1^ gentamicin did not have any significant impact on the biomass of *K. pneumoniae* and *E. coli* in the biofilm. Similarly, a 15 min treatment model C-5 with 4 mg ml^-1^ gentamicin did not significantly impact the *E. coli* and *E. faecalis* biomass in the biofilm. *K. pneumoniae* biomass slightly decreased from 1.5 × 10^8^ CFU cm^-2^ to 3.8 × 10^7^ CFU cm^-2^ while *P. aeruginosa* biomass decreased from 3.5 × 10^8^ CFU cm^-2^ to 2.1 × 10^7^ CFU cm^-2^.

**Figure 5.**
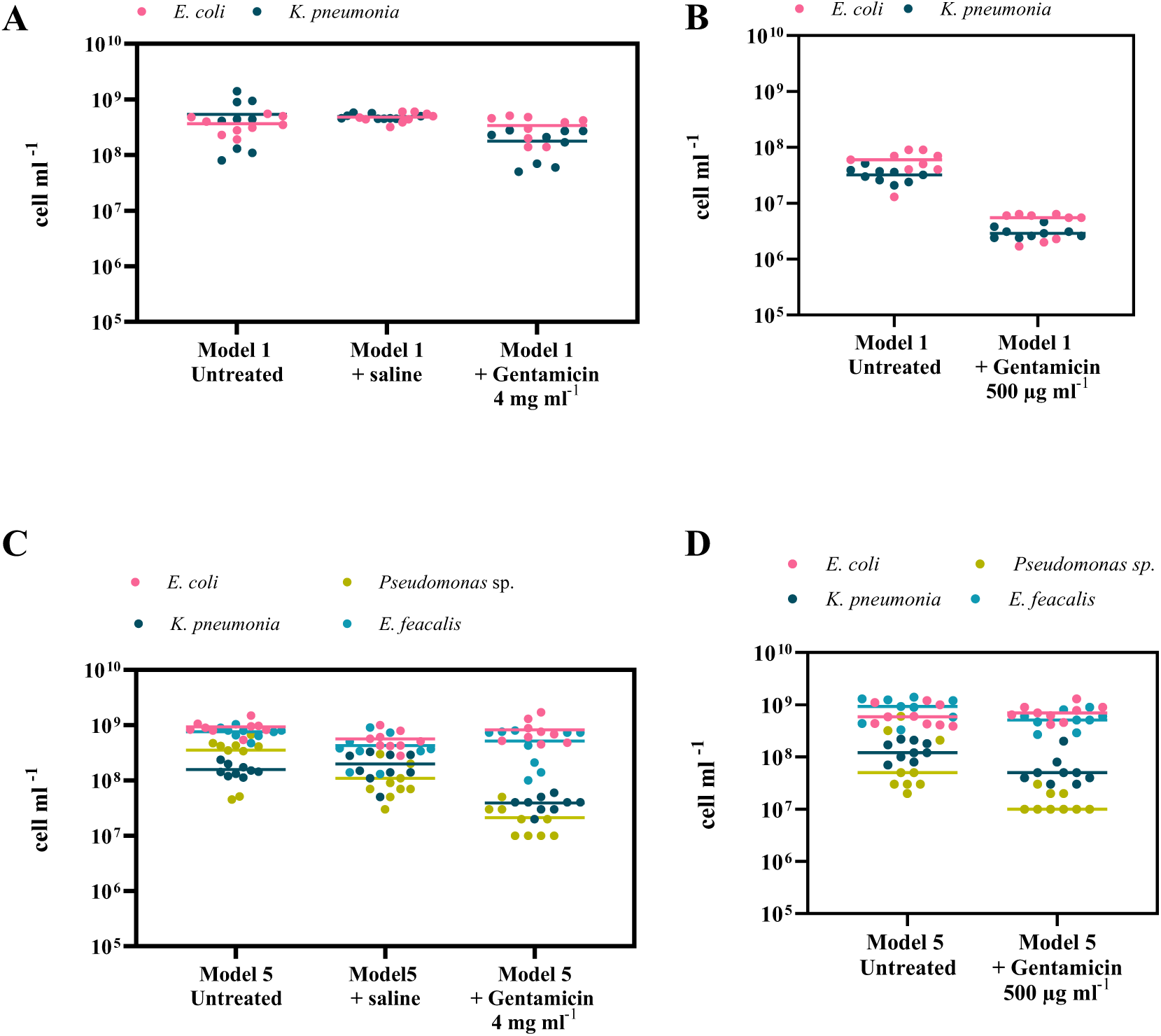
CFU counts on CHROMagar™ Orientation plates was used to determine the amount of. *K. pneumoniae, E. faecalis, P. aeruginosa* and *E. coli* in biofilms remaining on the catheters after wash based on the colour of each colony. A) Model C-1 and C) Model C-5 were exposed to 4 mg ml^-1^ gentamicin for 15 min. B) Model C-1 and D) Model C-5 were exposed to 500 µg ml ^-1^ gentamicin for 24 h. All experiments were repeated three times separately with three replicates in each independent experiment.

Furthermore, model C-1 multispecies biofilm was treated with this 500 µg ml ^-1^ of gentamicin for 24 h, which slightly decreased the biofilm biomass of *E. coli* from 6 × 10^7^ CFU cm^-2^ to 4.6 × 10^6^ CFU cm^-2^ and *K. pneumoniae* from 3.2 × 10^7^ CFU cm^-2^ to 3 × 10^6^ CFU cm^-2^. After 24 h exposure to 500 µg ml^-1^ gentamicin, model C-5 multispecies biofilm did not show any significant changes in the biomass of *E. coli* and *E. faecalis.* However, the biomass of *K. pneumoniae* and *P. aeruginosa* were slightly reduced by 0.5 and 1 log, respectively.

## Discussion

Biofilms on urinary catheters often present as complex, multispecies communities that can complicate treatment and management of catheter-associated infections (CAUTIs) ^26^. One of the primary challenges in creating a multispecies bacterial model is accurately replicating the microbial isolates that inhabit the human body and understanding the relationships among these organisms. To address this challenge, we took a novel approach that differed from previous studies that typically select isolates and artificially mix them in the lab. Instead, we started with pre-existing biofilm communities extracted from SCI patients who were asymptomatic and initially colonised by 5 to 10 species (Table 1). This allowed us to stabilize and cultivate this community under controlled laboratory conditions which resulted in final models of 2 to 4 species (Figure 1). This method intended to maintain a more natural relationships among the bacteria, leading to a more representative model for testing.

There was genetic diversity amongst strains of the four prominent species found across different models but there were no changes in the dominant strains in each *in vitro* model between passages (Figure 2). Analysis of *P. aeruginosa* isolates with two phenotypes in pigment production revealed several non-synonymous SNPs (Table 3). None of the genes carrying SNPs were directly associated with biosynthesis of pigments. However, regulatory genes and metabolic pathways in bacteria are often interconnected. For example, sensor protein RstB, part of the two-component RstA/RstB signal transduction systems, positively regulates the efflux pump MexEF-OprN, enzymes involved in anaerobic nitrate respiration, and pyoverdine biosynthesis ^27^.

The genetic diversity of our multispecies biofilm provides a more realistic test of treatments for bacterial eradication in clinical setting. Pathogens associated with CAUTIs often exhibit higher antibiotic resistance compared to those in non-catheter UTIs ^28^. This is on top of the inherent resistance characteristics of biofilms. Our data highlighted a concerning prevalence of antibiotic resistance genes across the biofilm communities. *S. agalactiae* had the lowest number matches of antibiotic resistance genes in all the databases used here, with only three identified *tetM* (tetracycline), *ErmB* (erythromycin) and *mprF* (cationic peptides), which was surprising given that it displayed resistance to four antibiotics including nalidixic acid and sulphametoxazole and intermediate resistance to seven in the disk inhibition assay. In contrast, *E. coli* from model C-1, which exhibited resistance to only one antibiotic and intermediate resistance to three in the disk inhibition assay, had 43 matches in the CARD database. Similarly, *K. pneumoniae* from model C-1 showed only intermediate resistance to only three antibiotics but had 22 matches in CARD. Model C-5 demonstrated a concerning level of resistance in the disk inhibition assay, which was reflected in the number of matches each species had in the NCBI and CARD databases. Specifically, the *E. faecalis* strain 5-3 exhibited the highest resistance among all tested isolates, displaying resistance to 15 antibiotics and intermediate resistance to two others. However, CARD analysis only identified a total of 10 matches for this strain. *E. coli* from model C-5 had 45 matches, while *K. pneumoniae* had 25 matches. *P. aeruginosa* also showed significant resistance, with 47 matches in CARD, and no differences were observed between the individual isolates 5-6 and 5-10. Many of the matches found in CARDS were those of efflux pumps which can act against many different antibiotics (Supplementary Data 2). These results underscore the complexity of the resistance mechanisms present within the microbial communities studied, revealing a diverse array of resistance genes and mechanisms. By identifying these genes, we can better understand the potential pathways through which microorganisms acquire and express resistance, ultimately informing strategies to combat AMR.

Gentamicin, specifically as an intravesical instillation, is a treatment for CAUTI in patients with neurogenic bladders and is recommended based on studies in which participants who underwent the antimicrobial instillation reported a good response with a reduction of symptomatic CAUTI ^29,30^. In many other cases, gentamicin is used as IV/IM, especially if sepsis is a concern.

The simplicity and reproducibility static multi well plates with biofilm growing on catheter pieces fixed to the lid of the plate as pegs allowed us to test all single isolates and mixed biofilm models. Drastic increase was observed in gentamicin resistance of all tested species but *K. pneumoniae* strain 1-5, when grown in multi-species. Also *S. agalacticae* could not be consistently observed in the mixed species biofilm possibly due to it’s very low percentage of the population. It is important to note that in both single or mixed scenarios, the bacteria had to compete for growth on the catheter. Interestingly, in all cases, there was higher total biomass when grown in mixed species biofilms compared to single species conditions.

Th architecture, structure and arrangement of bacteria within biofilm community is heavily implicated in antibiotic resistance ^31^. The formation of multispecies biofilm under flow conditions allowed us to simulate the dynamic conditions typically encountered in clinical settings, providing a more accurate representation of how these biofilms respond to antibiotic treatment. By maintaining two treatment scenarios: short 15 min intravesical treatment with high concentration versus a longer 24 h at lower concentration excreted in urine within the first 24 h after IV/IM administration, we aimed to create an environment conducive to biofilm formation and evaluate the effectiveness of gentamicin in this context. In all of these experiments, every species in both models demonstrated resistance to gentamicin, and the biofilms continued to persist after treatment, showing only minor changes in population percentage and abundance.

Overall, this work has simulated simplified clinical polymicrobial biofilm environments, in order to improve strategies for managing and mitigating microbial infections caused by biofilms. The comparative analysis of these polymicrobial communities as well as the development of a multi-species biofilm models for laboratory use have yielded significant insights into genetic diversity of bacteria and prevalence of antibiotic resistance mechanisms. Future efforts to develop antibiofilm strategies need to consider the intricate nature of polymicrobial biofilms, including their inherent resilience and resistance to treatment. It’s also essential to test these approaches in appropriate biofilm models to ensure their effectiveness.

## Methods

### Participant’s natural biofilm community

The samples for this project were obtained as part of a multi-centre observational longitudinal study, 3PU (ACTRN12622000613707). Biofilm collection was performed as part of the patient’s routine care or catheter change due to suspected UTI by their care team and only samples that would otherwise be discarded were collected over 18 months. The used catheters were explanted by trained health personnel. A 5 cm section of the bladder end of the catheter was cut using sterile scissors and collected in a sterile container containing 5 ml of sterile saline. After the new catheter were installed, fresh catch urine was collected from newly installed catheter in a sterile container. Both containers were transported to the research team for analysis. Biofilms were collected by cutting the catheter length wise and then physically detaching the biofilm using cell scraper. The catheter pieces were then submerged in saline and cells were extracted by vortexing briefly, sonicating in the ultrasonic cleaner, Power Sonic 420 (Thermoline) on medium for 1 min, followed by vortexing. Here, 1 ml of catheter cell suspension was centrifuged at 5000 g for 5 min. The supernatants were discarded and the cell pellets from the catheter were stored at -20°C for DNA extraction. The remaining catheter cell suspensions were centrifuged at 5000 g for 5 min and cell pellets were frozen at -80°C in glycerol.

### Establishing *in vitro* urinary catheter associated multi-species biofilms

Mixed species bacterial culture inputs were prepared using six biofilm samples selected at random from a pool of routine catheter changes stored in 25% glycerol. Samples were inoculated in complete composite Synthetic Human Urine (ccSHU) consisting of 100 mM NaCl, 17.0 mM Na_2_SO_4_, 280 mM Urea, 38.0 mM KCl, 4.0 mM CaCl_2_, 9.0 mM Creatinine, 3.4 mM Na_3_C_6_H_5_O_7_, 20.0 mM NH_4_Cl, 3.2 mM MgSO_4_, 0.18 mM Na_2_C_2_O_4_, 3.6 mM NaH_2_PO_4_, 6.5 mM Na_2_HPO_4_, 16.0 mM KH_2_PO_4_, 0.6 mM C_5_H_4_N_4_O_3_, 13.5 mM NaHCO_3_, 13.2 mM MgCl_2_, 1.1 mM C_3_H_6_O_3_, 0.005 mM FeSO_4_ supplemented with 0.1% (v/v) casamino acids, 0.2% (v/v) yeast extract ^19^, 0.5% Glucose and 0.5% Foetal calf serum (FBS). The pH was adjusted to 6.2 and sterilized by filtration through a 0.2 μm filter before use. Samples were incubated for 48 h at 37°C without shaking.

Releen Male Inline Catheters 18G, 10 cc, Silicone 40 cm (Coloplast) were aseptically cut into 10 cm lengths pieces. All catheter pieces were coated in 500 μg mL^-1^ Fibrinogen from human plasma (Sigma) overnight at 4°C. Masterflex^®^ peristaltic pumps (Cole-Parmer) with Masterflex^®^ L/S^®^ Multichannel Cartridge Pump Head with Reduced Pulsation for Microbore 2-Stop Tubing, 12 Channel, 8 Roller was used for flow delivery into catheters. All media flasks, connectors, pump and waste lines were autoclaved prior to use except for Masterflex^®^ L/S^®^ 2-Stop Precision Pump Tubing, Platinum-Cured Silicone which were sterilized using 0.25% Bleach solution for 1 h before being washed for 1 h using sterile distilled water. Pump lines were placed in inflow bottles containing ccSHU and then connected to Masterflex tubing, using straight connectors. The Masterflex tubing ran through the cartridge pump that had a steady flow rate of 50 μL min^-1^. This was connected using straight connectors to another segment of pump line that attached on to the 10 cm length piece of catheter and were connected to waste lines ending in waste flasks (Figure 6). Initially, ccSHU was allowed to run for 2 h to prime the system. Each sample of mixed species bacterial culture input was diluted to OD_600_=0.001 in 1 mL of ccSHU. The flow was paused and each end of the catheter was blocked to minimise the chance of backflow. Each culture input was loaded into a 1 mL syringe attached to a 26G 1/2 needle and directly injected into the catheter piece and sealed using silicone. To allow cell attachment, the flow was restarted 2 h after injection. The system was then allowed to run with flow of 50 μl min^-1^ for 4 d at 37°C, after which the inflow medium was replaced with saline and allowed to run for 2 h to remove planktonic bacteria. The catheter section was then removed and the extra liquid was drained. Each section was sliced horizontally in half using a sterile scalpel blade. The internal biofilm was recovered using a cell scraper and the pieces were submerged in saline and cells were extracted by vortexing briefly, sonicating in the ultrasonic cleaner, Power Sonic 420 (Thermoline) on medium for 1 min and then vortexing again. A portion of the extracted biofilm was kept at -20°C for total DNA extraction. A portion of the extracted biofilm was plated on CHROMagar™ Orientation plates to monitor the diversity of the biofilms based on chromogenic appearance of the colonies. Isolates from these plates were stored in 25% glycerol at -80°C and used as bacterial culture input for the next generation of biofilm flow (G1-G3). This process was repeated three times.

**Figure 6.**
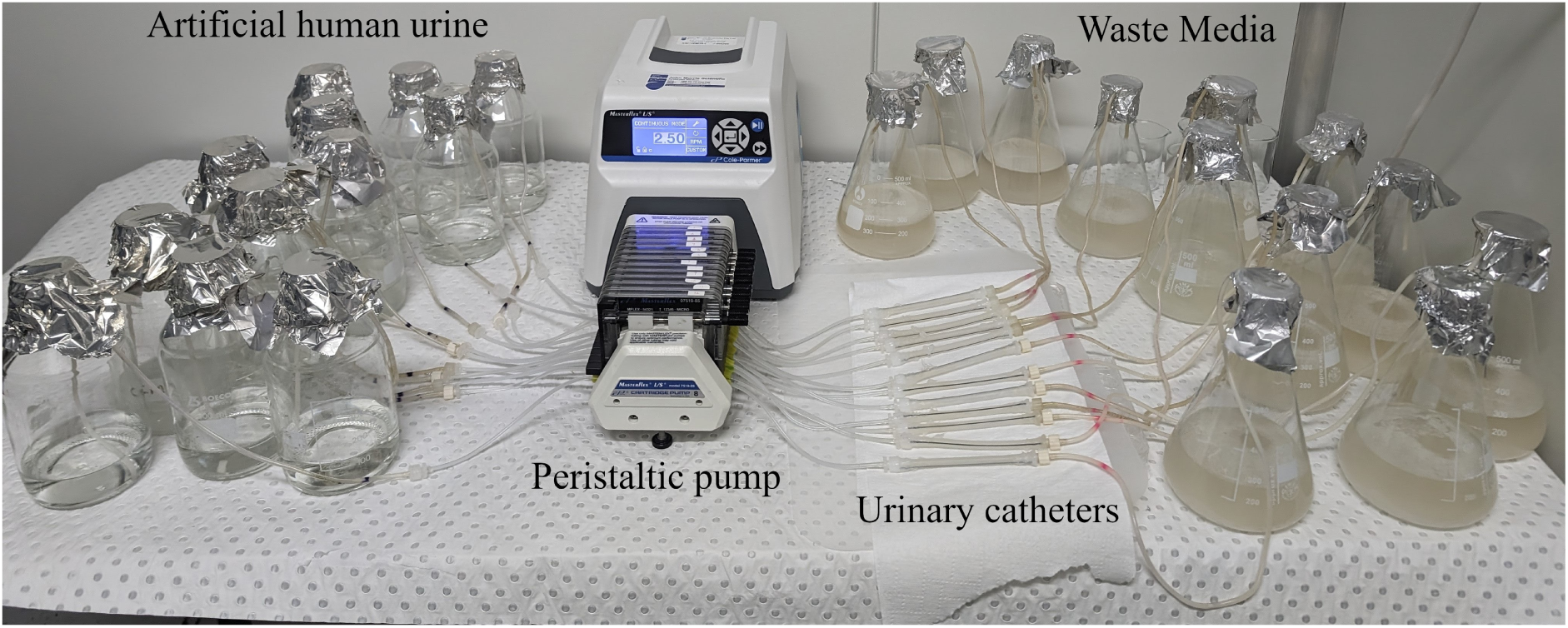
Photograph of the catheter flow biofilm model.

### Whole Genome Sequencing

Genomic DNA was extracted from biofilm samples using the DNeasy PowerSoil Pro Kit (QIAGEN) according to the manufacturer’s instructions. The quality and quantity of the isolated DNA were determined using a NanoDrop spectrophotometer (Thermo Fisher Scientific). The DNA samples were resuspended in TE buffer and stored at -20°C before further analysis. Genomic library preparation for all short-read sequencing was using the Hackflex library preparation method ^32^ and sequenced on NovaSeq PE150. The reads ranged from 81.5 - 4 .2 M reads. Samples explanted from participants were heavily contaminated with human genomic DNA. An Miseq sequencer was used to estimate the amount of human gDNA contamination and therefore, heavily contaminated samples were adjusted and sequenced deeper while samples that were from pure bacterial culture were shallow sequenced.

The G4 Input of models C-1 and C-5 mixed species bacterial culture was produced by inoculation of the of second generation (G3) biofilms output and incubation for 48 h at 37°C static. Genomic DNA was extracted using MagAttract HMW DNA Kit (QIAGEN) according to the manufacturer’s instructions. The DNA was then quantified using Qubit 3.0 fluorometer and NanoDrop One. Agilent 4200 TapeStation system and Agilent Femto Pulse systems were used for analysis of high-molecular-weight fragment size. DNALigation Sequencing Kit V14 (SQK-LSK114) and its recommended protocol for making sequencing libraries was used. Each library was then run on one PromethION Flow Cell separately. Nanopore sequencing signals were processed using MinKNOW 22.12.5 and base-called using Guppy Guppy 6.4.6+ae70e8f on the PromethION tower using the GPUs. This resulted in a total of 99,064,035,191 bp from which 78,941,083,703 bp (79.69%) passed with quality ≥Q9 and N50 of 6,432 bp for model C-1 and total bases 61,720,670,275 bp from which 48,685,922,888 bp (78.88 %) passed with quality ≥Q9 and N50 6,230 bp for model C-5.

The quality of the pair-end short reads was checked with FastQC ^33^, trimming and adaptor cut was performed by fastp set at default ^34,35^. Nanopore long reads were trimmed based on quality (in priority ≥90%) and length (12 kbp) and adaptor cut was performed by Fitlong ^36^.

### Metagenomic sequencing, and recovery and assessment of microbial populations

Short reads were used to determine the taxonomic classification using Kraken2-2.1.3 ^37^. The ready to use standard plus Refeq protozoa & fungi database (77 GB) available from the Langmead lab website (https://benlangmead.github.io/aws-indexes/k2; 09-10-2023 version) was used. Minimum hit groups and confidence were set to 4 and 0.3, respectively. Followed by Braken V2.9 for a taxonomic abundance estimation and Pavian for visualisation ^38,39^. All samples were heavily contaminated with human DNA. Here, the percentages are reported after filtering out human reads using Pavian Homo Sapiens option as well as the removal of all phage reads.

Genomes of the single isolates were assembled using SPAdes v.3.15.5 genome assembler ^40^. Aviary’s ^41^ Metagenome-assembled genomes (MAGs) recovery workflow was used to co-assemble the short and long reads alongside MAG recovery, binning and annotation. Various different tools were used, including NanoPack and NanoPack2 ^42,43^, fastp ^34,35^, Flye ^44^, Racon ^45^, Pilon ^46^, metaSPAdes ^47^, Unicycler ^48^, Minimap2 ^49^, samstools ^50^, CONCOCT ^51^, VAMB ^52^, MetaBAT 1 and MetaBAT 2 ^53,54^, DAS Tool ^55^, MaxBin 2.0 ^56^, SemiBin 2 ^57^, Rosella ^58^, CheckM and CheckM2 ^59,60^, eggNOG mapper 2 ^61^ and GTDB-Tk 2 ^62^. Briefly, hybrid assembly of short and long reads was performed using workflow “assemble.” The assemblies were manually checked using MetaQUAST ^63^ and Bandage ^64^. Genomes of each community were then recovered using workflow “recover.” Quality of MAGs was checked using CheckM2 version 1.0.2. Taxonomy information of MAGs was determined using GTDB-Tk 2.3.3 and taxonomy of the single isolates was determined using GTDB-Tk 2.4.0. ABRicate V1.0.1 (https://github.com/tseemann/ABRicate) all databases update on 04/11/2023 was used to screen for AMR genes with the NCBI AMRFinderPlus ^65^, The Comprehensive Antibiotic Resistance Database (CARD) ^66^, EcOH.

Metaphlan 4.1.1 ^67^and Strainphlan 4.1 ^68^ were used to track individual strains across samples and determine whether there was strain specific divergence or the presence of a multi-strain species complex. The MetaPhlAn mpa_vJun23_CHOCOPhlAnSGB_202307 was used to map reads to marker sequences.

The breseq v0.38.3 pipeline ^69^ was used to compare the single isolates of same species from each model that showed different phenotype. Briefly, prokka 1.14.6 ^70^ was used to annotate each genome. The annotated isolates were used as reference by breseq pipeline to identify for mutations and *gdtools* program were used for comparison between samples ^71^.

### Antibacterial testing

Generation 4 (G4) of Models C-1 and C-5, were used for the antibacterial testing. Single bacterial isolates were selected by using colony morphology and colour on CHROMagar™ Orientation plates.

Single isolates were grown overnight in Brain Heart Infusion (BHI) (BD BACTO™) broth at 37°C, 200 rpm. The day of the test, cultures were diluted and regrown to mid-log phase to absorbance 0.5 at 600_nm_. Antibiotic discs were placed on Mueller-Hinton Agar (MHA) plates using antimicrobial susceptibility disc dispenser (Oxoid™) followed by overnight incubation for 18 h at 37°C. The antibiotic disks (Oxoid™) used in this study included: Amikacin 30 µg, Ampicillin Sulbactam 20 µg, Ampicillin 25 µg, Cefotaxime 30 µg, Ceftazidime 30 µg, Ceftriaxone 30 µg, Chloramphenicol 30 µg, Ciprofloxacin 5 µg, Gentamicin 10 µg, Imipenem 10 µg, Kanamycin 30 µg, Meropenem 10 µg, Nalidixic Acid 30 µg, Neomycin 30 µg, Netilmicin 30 µg, Rifampicin 30 µg, Spectinomycin 25 µg, Streptomycin 25 µg, Sulfamethoxazole 100 µg, Tetracycline 30 µg, Tobramycin 10 µg, Trimethoprim 5 µg ^72^.

The antimicrobial susceptibility of planktonic cells was performed using a resazurin-based microdilution assay ^73^. Briefly, overnight cultures of each strain or the established mixed model G4 input were grown overnight in ccSHU and adjusted to an OD_600_ = 1.0. Each strain was then diluted 1000-fold in ccSHU. Each sample was then inoculated in a 96 well microtiter plate with different concentrations of the selected antibiotics. Each plate was incubated at 37°C, static for 20 h. After the incubation period, 5 µl of 1 mg mL^-1^ Resazurin (Sigma) was added to each well incubated at 37°C for 1 h. Results were quantified using fluorescence spectrophotometer excitation at 575 nm and an emission at 585 nm using Spark^®^ microplate reader (Tecan). A positive control of isolates in ccSHU with no antibiotics and a negative control of antibiotics without bacteria was used in this assay.

Releen Male Inline Catheters 18G, 10 cc, Silicone 40 cm (Coloplast) were aseptically cut into 15 mm lengths pieces. The inflation line was removed using a sterile scalpel blade and each piece was fixed to the lid of a 48 well plate (Figure 7). Overnight cultures of each isolate were grown in BHI broth at 37°C, 200 rpm. Mixed biofilm samples were grown in ccSHU for 48 h at 37°C without shaking. Each culture was then washed in SHU and adjusted to an OD_600_ = 0.1. The cultures were diluted 100-fold in ccSHU and inoculated in 48 well plates with modified lids. Each plate was incubated at 37°c, static for 48 h and the medium was refreshed every 12 h. Fresh ccSHU with or without gentamicin for 500 µg ml^-1^ was subsequently added for a further 24 h and the medium was again refreshed after a further 12 h. The catheter pieces were then washed in saline, 9 g L^-1^ NaCl_2_ before detachment of each catheter from plate. The biofilm was scrapped and the pieces were submerged in saline and cells were extracted by vortexing and ultrasonic cleaner. The bacterial biomass was then enumerated using CFU on BHI agar for single isolates and CHROMagar™ Orientation for multispecies samples.

**Figure 7.**
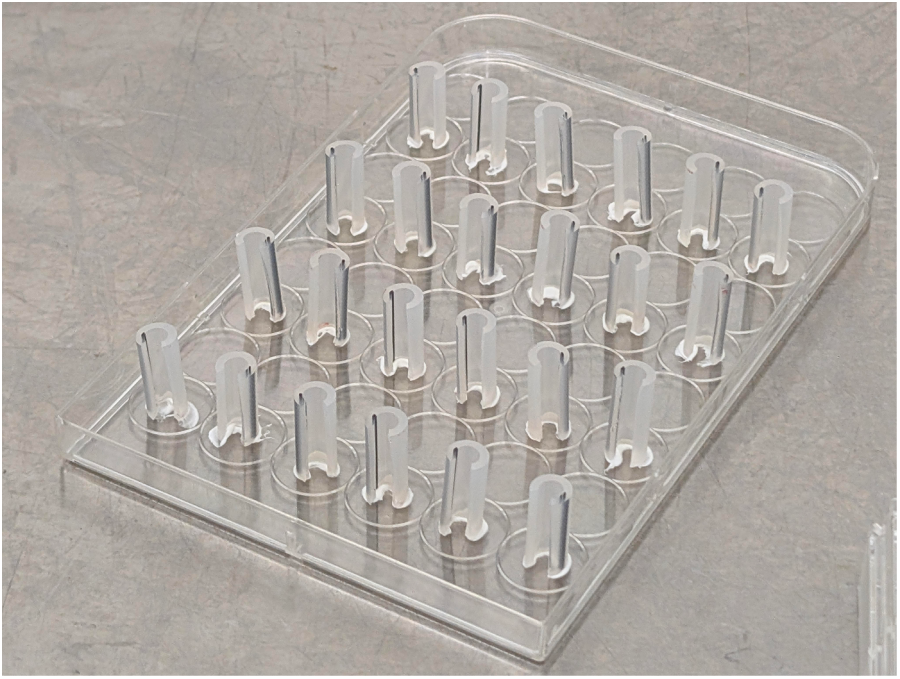
Photograph of catheter pieces (1.5 cm) Releen Male Inline Catheters 18G, 10 cc, Silicone 40 cm (Coloplast) with inflation lines removed fixed as pegs to lid of a multi well plate.

To test multi-species biofilm antimicrobial susceptibility under flow conditions, 4 d biofilms were established on 10 cm length catheter pieces. Each condition was tested in 3 separate lines for each repeat and the experiments were repeated as 3 independent experiments. Two timepoints and concentrations were used to test antibiotic resistance. Doses of 80 mg of gentamicin per 20 ml of saline instillation for 15 min are frequently used as a bladder irrigation technique ^74^. In order to test the effectiveness of the antibiotic without the help of the physical force of the wash, saline or gentamicin 4 mg ml^-1^ were injected gently using a syringe to the lines containing 4 d old biofilms. Upmost care was used to inject very gently to eliminate the disruption of biofilm via physical force. After 15 min, the lines were washed for 2 h with saline only and all of the liquid was gently drained from them. The catheter section was then removed and sliced horizontally in half using sterile scalpel blade. The internal biofilm was scrapped and the pieces were submerged in saline and cells were extracted by vortexing and ultrasonic cleaner. The bacterial biomass was then enumerated using CFU on CHROMagar™ Orientation. Furthermore, the Input bottles for the rest of the lines were switched to fresh ccSHU with or without gentamicin for 500 µg ml^-1^ for another 24 h under the same flow conditions. After this all lines were washed for 2 h with saline only, and all the liquid was drained gently from them. The remaining attached cells were then removed as described above and quantified.

### Statistical analysis

Statistical analysis was performed using GraphPad Prism version 10.0.2 for Windows, GraphPad Software, La Jolla California, USA (www.graphpad.com). Data that did not follow Gaussian distribution as determined by analysing the frequency distribution graphs was natural log transformed. Two-tailed Student’s t-tests were used to compare means between experimental samples and controls. For experiments including multiple samples, one-way or two-way ANOVAs were used for the analysis, and Sidak’s or Dunnett’s Multiple ComparisonTest provided the post-hoc comparisons of means when appropriate. Data organization, visualization and formatting were conducted using the R package “tidyverse”, “ggplot2” and “ggtree” ^75,76^.

### Ethics approval and consent to participate

The samples for this project were obtained as part of a multi-centre observational longitudinal study 3PU (ACTRN12622000613707). The ethics approval for all study protocols were obtained from the Northern Sydney Local Health District Human Research Ethics Committee (HREC) 2021/ETH00329 as well as Site Specific Assessment (SSA) 2021/STE00542 by Royal North Shore Hospital governance, 2021/STE00541 by Prince of Wales Hospital governance and Royal Rehab governance dated 29/9/2022. Participants were adults, age ≥18 years, with stable SCI and stable neurogenic bladder management technique for at least 4 weeks before start of the study. Participants agreed to have their urinary system microbial community and the microbiome DNA from the catheter and urine to be extracted and stored long-term. Written informed consent was obtained from all participants and samples were de-identified by the assigning of an arbitrary subject number. Research Electronic Data Capture (REDCap) was used to capture demographic data as well as sample information.

## Supplementary Information

**Supplementary Data 1**: Details and statistic summary of MAG assemblies.

**Supplementary Data 2**: AMRs were detected in MAG against Comprehensive Antibiotic Resistance Database (CARD). Each AMR is categorized based on type and product and Bins.

## Author contributions

DM, SAR, IGD and BBL developed the concept and designed the study. PN and OM designed the participant data collection forms and questionaries. PN, JT, PC, GW, JP and BBL co-ordinate the recruitment of all participants, collection of the samples, and fortnightly telephone conversation. PN, JT, GEV, MMH and DL processed samples. PN, JT and KH extracted DNA from samples and performed sequencing. PN processed all the sequencing data, performed bioinformatics and statistical analyses. PN, JT and KH performed all the biofilm assays. PN wrote the manuscript with input from all authors. DM, SAR, IGD and BBL supervised the project. All authors provided valuable feedback to improve the final manuscript.

## Data availability

The datasets generated for this study can be found in online repositories. Raw reads of the single isolates and metagenomes are available at NCBI’s Sequence Read Archive (SRA) under Bioproject PRJNA1174239.

## Acknowledgements

We thank all the participants and care team who sincerely provided us the samples, especially clinical nurses Ruth Hamilton, Ruyi Yao and Jennifer Greenway. We also like to thank staff at Ferguson Lodge and staff of Fairfield West Clinic and Dr. Zahra Rassoly Obayd for their help with participant recruitment and sample collection.

## Consent for publication

Not applicable

