## Supplementary Data 1 for "A model, mixed-species urinary catheter biofilm derived from spinal cord injury patients": Supplementary Data 1.pdf

Supplementary Data 1:Details of MAG assemblies.

| | | | | | |
|---|---|---|---|---|---|
| | | | **Model 1** | | |
| **Bin Id** | 001_maxbin_bins | 002_maxbin_bins | 001_semibin_refined_bins | 008_concoct_bins | 003_rosella_refined_bins |
| **Total_Contigs** | 6 | 3 | 241 | 13 | 156 |
| **N50 (contigs)** | 5272598 | 5237876 | 6399 | 3363 | 794 |
| **Mean contig length (bp)** | 933193 | 1765743 | 4807 | 3182 | 531 |
| **Longest contig (bp)** | 5272598 | 5237876 | 28499 | 14174 | 1477 |
| **Genome size (bp)** | 5599158 | 5297229 | 1760820 | 50954 | 85086 |
| **Completeness /contamination (CheckM1)** | 100 / 0.04 | 99.97 / 0.08 | 90.91 / 062 | 2.59 / 0 | 3.61 / 0 |
| **Completeness/contamination (CheckM2)** | 100 / 0.33 | 100 / 1.31 | 92.49 / 2.69 | 3.74 / 0 | 5.07 / 0 |
| **classification** | *Klebsiella pneumoniae* | *Escherichia coli* | *Streptococcus agalactia* | Unclassified | Unclassified |
| **fastani_reference** | GCF_000742135.1 | GCF_003697165.2 | GCF_000186445.1 | | |
| **fastani_reference_radius** | 95.239 | 95 | 95 | | |
| **fastani_ani** | 98.91 | 98.65 | 98.72 | | |
| **fastani_af** | 0.881 | 0.909 | 0.917 | | |
| | | | **Model 5** | | |
| **Bin Id** | 002_maxbin_bins | 003_maxbin_bins | 004_maxbin_bins | 005_metabat2_refined_bins | 001_sub_maxbin_bins |
| **Total_Contigs** | 6 | 8 | 47 | 1 | 8 |
| **N50 (contigs)** | 2951105 | 2126134 | 307141 | 5080905 | 13597 |
| **Mean contig length (bp)** | 518527 | 696425 | 153111 | 5080905 | 14359 |
| **Longest contig (bp)** | 2951105 | 2412685 | 1083015 | 5080905 | 44870 |
| **Genome size (bp)** | 3111164 | 5571401 | 7196254 | 5080905 | 114876 |
| **Completeness /contamination (CheckM1)** | 99.63 / 0.37 | 100 / 0.56 | 96.79 / 0.45 | 99.97 / 0.39 | 0 / 0 |
| **Completeness/contamination (CheckM2)** | 100 / 0.39 | 100 / 0.75 | 96.35 / 0.22 | 100 / 0.85 | 4.52 / 0.17 |
| **classification** | *Enterococcus faecalis* | *Klebsiella pneumoniae* | *Pseudomonas aeruginos* | *Escherichia coli* | Unclassified |
| **fastani_reference** | GCF_000392875.1 | GCF_000742135.1 | GCF_001457615.1 | GCF_003697165.2 | |
| **fastani_reference_radius** | 95 | 95.239 | 95 | 95 | |
| **fastani_ani** | 98.65 | 99.02 | 99.27 | 98.67 | |
| **fastani_af** | 0.904 | 0.889 | 0.957 | 0.921 | |
